# MRI-derived brain age as a biomarker of ageing in rats: validation using a healthy lifestyle intervention

**DOI:** 10.1101/2021.04.19.440433

**Authors:** Irene Brusini, Eilidh MacNicol, Eugene Kim, Örjan Smedby, Chunliang Wang, Eric Westman, Mattia Veronese, Federico Turkheimer, Diana Cash

## Abstract

MRI data can be used as input to machine learning models to accurately predict brain age in healthy human subjects. A large difference between predicted and chronological brain age (the so-called BrainAGE score) has been associated with disease and neurodegeneration, indicating the potential utility of neuroimaging-based ageing biomarkers. So far, most brain age prediction studies have been carried out on humans. However, it is important for such a biomarker to be validated on laboratory animals too, in order to better account for specific environmental or genetic factors within a more controlled laboratory framework.

In this work, we developed a new algorithm for rat brain age prediction based on the combination of Gaussian process regression and a logistic regression classifier. The algorithm was trained on a cohort of 31 normal rats. High prediction accuracy was achieved using leave-one-out cross-validation (mean absolute error = 4.87 weeks, correlation between predicted and chronological age *r* = 0.92), supporting the validity and potential of the method.

Furthermore, the trained model was tested on two independent groups of 24 rats each: a new normal control group and a “healthy lifestyle” group that underwent long-term environmental enrichment and dietary restriction (EEDR) between 3 and 17 months of age. After fitting a linear mixed-effects model, the BrainAGE values were found to increase more slowly with chronological age in the EEDR group than in the controls (slope = 0.52 vs. 0.61; *p* = 0.015 for the interaction term). When survival analysis was performed with a Cox regression model, the BrainAGE score at 5 months of age had a significant prediction power (*p* = 0.03).

Our results demonstrate that BrainAGE, as computed by the proposed approach, is significantly modulated by EEDR intervention, hence it is a sensitive marker of biological ageing. These findings also support the potential of lifestyle-related prevention approaches to slow down the brain ageing process. Moreover, the results of the survival analysis further demonstrate that BrainAGE is indeed a predictor of ageing outcome.

## 1 Introduction

Due to the continuous improvement in the quality of life and healthcare, the world population is living longer, and the number of older individuals is growing remarkably (He et al., 2016). This longer life expectancy comes at a cost, i.e. increasing prevalence of diseases and functional decline that are associated with ageing (Denver and McClean, 2018). Ageing constitutes, for example, one of the major risk factors of developing dementia, a clinical syndrome affecting the brain whose symptoms include memory loss, language disturbances and a general impairment in daily activities (Burns and Iliffe, 2009). Moreover, age-related changes in peripheral body parts can also affect the brain itself, suggesting that good overall health is fundamental for maintaining a healthy brain (Cole and Franke, 2017). However, understanding and modelling how the brain ages is still a challenge, especially because this process is extremely heterogeneous across individuals. Age-related brain pathologies are indeed characterised by a very broad range of onset ages (Cole et al., 2017b).

It is now widely accepted that the age of the brain can differ from the person’s chronological age and that this is influenced by a variety of complex genetic and environmental factors (Lee and Sachdev, 2014; Lu et al., 2004; Peters, 2006; Teter and Finch, 2004; Rando and Chang, 2012; Cole et al., 2018). Despite the complexities, a better understanding of brain ageing and a link to neurodegeneration is essential to deal with the consequences of longer life expectancy (Franke and Gaser, 2019). If we can accurately assess brain age, we could use this as a biomarker of age- or disease-related pathologies to develop better treatments and improve the overall quality of ageing. To this end, research studies have begun to focus on the identification of reliable brain ageing biomarkers, which could be used to monitor age-related cognitive impairments, as well as detect neurodegenerative processes at their earliest stages (Cole et al., 2017b). Neuroimaging methods are ideally suited to such analyses due to their non-invasive nature, relatively wide accessibility and rapidly expanding number of publicly available data sets and software for brain image analysis.

To date, the best known brain age prediction studies involved the implementation of machine learning models with different magnetic resonance imaging (MRI) modalities as input, e.g. functional MRI (Dosenbach et al., 2010) or structural T1-weighted (T1w) MRI (Franke et al., 2010). The latter approach, similar to that used in our study, first pre-processes the raw T1w data using voxelbased morphometry (Ashburner and Friston, 2000), which includes tissue segmentation and spatial registration to a reference template. This preprocessing step allows extraction of biologically meaningful image features that relate to ageing—such as local grey matter (GM) concentration—and that are directly comparable across subjects. This is followed by a data dimensionality reduction step to prevent over-fitting and reduce the computational costs. Finally, a machine learning-based regressor is employed to model brain age from the processed MRI data. Recently, a growing number of more advanced deep learning-based approaches, including residual convolutional neural networks (Jónsson et al., 2019) and the Inception-ResNet-v2 framework (Bashyam et al., 2020), have been developed for the same purpose and obtained accurate brain age prediction results. However, such methods require large amounts of training data, and preclinical datasets of appropriate size are currently unavailable.

Prior studies have reassuringly confirmed that brain MRI can be used to predict both the chronological brain age in healthy subjects and a mismatch between biological and chronological age in clinical populations. In general, imaging-based brain age prediction models are trained on scans from healthy individuals and tested on new heterogeneous data from unseen subjects. If, during testing, the brain age is predicted to be *greater* than the subject’s chronological age, this could indicate the presence of a disease or neurodegeneration (Cole and Franke, 2017). If, on the other hand, brain age is predicted to be *lower* than the chronological age, this could reflect a favourable trend in an individual’s ageing process. The difference between predicted brain age and chronological age has been referred to as a *BrainAGE score* (Franke and Gaser, 2019), as defined in one of the first age prediction studies by Franke et al. (2010). Using such approach, BrainAGE has been confirmed a useful biomarker of abnormal brain ageing in patients with various neuropsychiatric disorders. Increased BrainAGE scores were found to be strongly associated with conditions such as epilepsy (Pardoe et al., 2017), traumatic brain injury (Cole et al., 2015), schizophrenia (Nenadićet al., 2017), Down’s syndrome (Cole et al., 2017a) and HIV (Cole et al., 2017c). Importantly, the predictive utility of a brain age biomarker was shown in studies of mild cognitive impairment, where individuals who converted to Alzheimer’s disease within 3 years had higher BrainAGE scores compared to those who remained disease-free (Gaser et al., 2013; Löwe et al., 2016). Increased BrainAGE scores could also be observed with peripheral pathological conditions, including diabetes (Franke et al., 2013) and mid-life obesity (Ronan et al., 2016). On the other hand, positive modifiers such as individual’s physical activity, as well as the number of years of education, were found to be associated with a decreased brain age (Steffener et al., 2016). Moreover, a study by Cole et al. (2018) found an association between predicted brain age and mortality risk.

The above-mentioned research strongly suggests that MRI-based brain age prediction is a promising new biomarker of ageing. The American Federation for Aging Research had outlined a series of criteria for ageing biomarkers (AFAR, from the *Infoaging Guides*, 2016 edition), one of which is that they should be applicable to both humans and laboratory animals, so that they can be extensively tested and validated preclinically before being fully accepted into a clinical framework (Johnson, 2006). In a previous work by Franke et al. (2016), two new species-specific adaptations of the BrainAGE model were tested on baboons and on rats. In both, the prediction model achieved accurate results, demonstrating the effectiveness in animal studies. In particular, the rat-specific BrainAGE model achieved a correlation of 0.95 between chronological and predicted age, with a mean absolute error (MAE) of 49 days. However, this model was only validated on a single cohort (using cross-validation), and its performance has not been tested on any experimental model in which genetic and environmental factors could be manipulated.

In this work, we developed a novel MRI-based algorithmic predictor of brain age in a preclinical laboratory setting on rats. We aimed not only to test its predictive ability and compare it to previous work, but also to investigate its sensitivity to ageing modulation intervention. On the methodological side, we introduce a novel algorithm for using T1w MRI data to predict brain age. Our strategy is based on the use of both Gaussian Process Regression (GPR) and a logistic regression (LR) classifier in order to minimise the prediction error in a training cohort of rat images used to fit the model. Subsequently, we tested the trained model on a separate cohort that included two groups of rats: a control group and an “active lifestyle” group that underwent environmental enrichment and dietary restriction (EEDR) between their early and late life (between approximately 3 to 17 months of age). Previous studies have robustly proved that environmental enrichment contributes towards long-term improvements in cognition and memory in both rodents and humans (Speisman et al., 2013; Hötting and Röder, 2013). These are believed to be a consequence of augmented brain plasticity, synaptic remodelling and neurogenesis, which are normally reduced in older age. Moreover, dietary restriction is also associated with increased neural plasticity and cognition, as well as neuroprotection in response to trauma and neurodegenerative disorders (Mattson, 2010; Martin et al., 2006). Thus, the present work also attempted to compare the BrainAGE scores of the control rats against the EEDR ones, in order to additionally investigate the effect of lifestyle modification on the ageing process within a controlled preclinical framework.

## 2 Materials and Methods

### 2.1 Animals

Male Sprague-Dawley rats were received from Charles River UK at 4.5 ± 0.5 weeks. All animal experiments were performed according to the UK Home Office Animals (Scientific Procedures) Act (1986) and approved by the Animal Welfare Ethical Review Body (AWERB) of King’s College London. The rats were divided into two cohorts: a *training cohort* of 31 normal ageing rats, and a test cohort—which will also be referred to as the *ageing cohort* —including 24 control subjects (i.e., which had a comparable lifestyle to the training cohort) and 24 EEDR rats. Environmental enrichment was obtained by using four different sets of toys (e.g., hammocks, hanging bells, and chewstick puzzles), which were changed weekly into the rats’ cages. Dietary restriction was achieved by removing food on three non-consecutive days every week for 24 hours, while on the remaining four days food could be accessed *ad libitum*. The control rats had *ad libitum* access to food every day and their cages were supplied with the standard enrichment only, that was wooden chewsticks and a cardboard tube. EEDR intervention commenced when the rats were 3 months old and immediately after the first scan session.

The rats were scanned in a maximum of four sessions, approximately at 3, 5, 11 and 17 months old, which correspond to adolescence, young adulthood, middle age and the start of senescence in humans (Quinn, 2005). All rats were scanned at the first session, but due to various factors, several rats were not scanned in some or all of the later sessions (see Table 1). Rats were excluded from scanning sessions if they became too large for the radiofrequency coil (above 850g) or for health and welfare reasons, such as age-related diabetes, arthritis and presence of tumours (N = 13). Finally, if a rat could not be socially-housed, it was excluded (N = 12) as the stress of social isolation (single housing) was expected to be a confound. One rat was excluded after data acquisition, due to the presence of a latent brain mass that impacted brain volume.

**Table 1:** Number of image samples for each scanning session and each of the analysed rat groups.

| Group | Number of animals |  |  |  |
| --- | --- | --- | --- | --- |
|  | Session 1 | Session 2 | Session 3 | Session 4 |
| Training cohort | 31 | 27 | 20 | 11 |
| Ageing cohort - controls | 24 | 24 | 17 | 10 |
| Ageing cohort - EEDR | 24 | 24 | 23 | 20 |

### 2.2 MRI image acquisition

For scanning, the rats were anaesthetised with 5% isoflurane in a 30:70 mixture of oxygen in air, with a flow of approximately 1 litre per minute. Isoflurane was then reduced to 2.5% during scanning. The scan bed included a built-in heating system using hot water, which was supplemented with a tube supplying thermostatically-controlled hot air to maintain body temperature at 37 ± 1°C. A rectal thermometer, a pulse-oximeter and a respiration sensor (made by Small Animal Instruments, Inc., NY, USA) were used to monitor the physiology. Each rat was scanned, head prone, in a 9.4 T Bruker Biospec MR scanner with an 86 mm volume transmission coil and a four-channel array receiver coil placed on the superior head surface, and a transmit/receiver ASL coil inferior to the rat’s neck to transmit labelling pulses for cerebral blood flow measurements (results not reported here) using scanning protocols implemented on Paravision 6.0.1 (Bruker Corp., Ettlingen, Germany).

High resolution 3D anatomical brain images were obtained by using an MP2RAGE sequence with the following parameters: repetition time (TR) = 9000 ms; inversion times (TIs) = 900, 3500 ms; flip angle = 7°, 9°; echo time (TE) = 2.695 ms; echo TR = 7.025 ms; matrix = 160 *×* 160 × 128; and 0.19 mm isotropic voxel size. A 3D ultra-short echo (UTE) reference scan was also acquired by setting: TE = 8 s, TR = 3.75 ms, flip angle = 4°, matrix = 128 × 128 × 128, and 0.45 mm isotropic voxel size.

The complex images from each coil were combined using the UTE reference scan and applying the COMPOSER method (Robinson et al., 2017), as implemented by Quantitative Imaging Tools v2.0.2 (Wood et al., 2016). The combined images were then used as input to `qimp2rage` in order to produce both a T1 map and a T1w image. The latter was then reoriented to RAS orientation.

### 2.3 Frailty index (FI)

Frailty is characterised by an increasing likelihood of poor health outcomes and the likelihood of frailty increases with age. Healthy ageing is associated with minimal frailty, so quantifying the degree of frailty is essential for describing heterogeneity in health outcomes in ageing studies. The rats’ FI was scored by a single researcher (author EM) at most two days before their session 3 and 4 scans, with a truncated protocol based on the criteria described by Yorke et al. (2017). Briefly, the rats were placed in a clean cage and a 25 point FI assessment was undertaken measuring clinical signs and deficits across a range of systems including integument, musculoskeletal system, ocular/nasal systems, digestive/urogenital systems, respiratory system, as well as assessment of discomfort and body weight. The protocol notably deviated from Yorke et al. (2017)’s FI protocol in the hearing loss test: we measured the presence of a startle reflex in response to clicking an unloaded stapler, out of sight and approximately 15 cm from the subject three times with 30 seconds between each click. Rats that startled at all three clicks were given one point, while a score of 0.5 was given to rats that startled to a minimum of one click. Rats that did not startle were assigned zero.

Group FI scores are reported as the mean ± standard deviation. The effect of session and group on FI was tested by fitting a mixed-effects linear model using GraphPad Prism v9.1.0. Posthoc tests between groups used Sidak’s multiple comparison test, and *p* < 0.05 was considered statistically significant.

### 2.4 Image preprocessing

The T1w image of each animal was skull-stripped using a modified implementation of `artsBrainExtraction` (MacNicol et al., 2020), an atlas-based algorithm for rodent brain extraction that involves registering individual subjects to a reference template with a predefined brain mask. We employed the 11-month-old rat brain template generated by MacNicol et al. (2021) as the reference template, as it constitutes a “middle age” reference that minimises the differences between all potential subjects and the template. The template image has dimensions of 160 × 160 × 128, with 0.19 mm isotropic voxel size. The reference atlas from the same study was also used to extract three tissue probability maps (TPMs)—for GM, white matter (WM) and cerebrospinal fluid (CSF)—for each subject by employing the ANTs `Atropos` tool (Avants et al., 2011) with consistent parameters for all subjects.

Finally, the extracted TPMs were modulated by multiplying them by the Jacobian determinants of the transforms obtained from the previous template-registration step. In this way, for each subject, modulated GM, WM and CSF TPMs—all defined in template space—were available to be used as input to the age prediction model.

### 2.5 Brain age prediction model

The proposed pipeline for rat brain age prediction was implemented in Python and optimised using a leave-one-out cross-validation approach with the available training cohort. The methodology that we followed can be divided into three main steps, presented in the subsections below.

#### 2.5.1 Input data preparation

For every subject, each modulated TPM was loaded and flattened into a one-dimensional vector. We then investigated different alternatives as possible inputs for the age prediction model. First, we tested the following inputs: (1) either the modulated GM or WM probability map as a unique input for the model; (2) the concatenation of both modulated GM and WM probability maps, but discarding the CSF; (3) the concatenation of all the three modulated TPMs (GM, WM, CSF). Later, we decided to generate additional TPMs not only using the 11 months reference template (as described in Section 2.4), but also using all other available age-specific templates (i.e. for 3, 5 and 17 months). We then investigated the same input configurations described previously (i.e. GM only, WM only, GM+WM, GM+WM+CSF), but now additionally concatenating all the respective TPMs obtained from the four available templates. This last approach requires a longer and more intensive preprocessing of the input images, but we believed that it was worth investigating whether a combination of the different templates could provide higher prediction accuracy. However, as further explained later in the Results section and in the Supplementary Material, we ultimately chose to only use the three modulated TPMs (GM, WM, CSF) from the 11 months template as input to the model. In this way, for each subject, a final input vector of size 1 × 9 830 400 was obtained.

Once all the input vectors (one for each rat and each scanning session in the training set) were generated, they underwent principal component analysis in order to reduce data dimensionality. Only the first 77 principal components (PCs) were kept, which preserved up to 95% of the total variance. In this way, the size of each input sample was transformed to 77 features, which correspond to the coefficients of the selected PCs.

These input samples were used to train both the GPR and the LR models (described in the next sections) in a leave-one-out cross-validation fashion. At each iteration, a different rat was selected as part of the validation set, while all remaining 30 rats were employed to train the models. Both models were implemented and fitted using the Scikit-learn v.0.23.2 Python library (Pedregosa et al., 2011).

#### 2.5.2 Gaussian process regression (GPR) model

As a baseline model for the present work, we employed a GPR model with a linear kernel. This model has already been used for predicting human brain age and showed accurate results (Cole et al., 2015). As input for the model, we used the concatenated TPMs projected onto the PC space. Moreover, the chronological ages (i.e., the outputs to be predicted) were provided as a column-vector containing all ages expressed in weeks. This unit of time was the most suitable choice for our data set, since the rats’ birth dates were provided by the animal supplier with an uncertainty of ± 4 days.

#### 2.5.3 Integration of linear regression (LR) predictions

In addition to the above-described model, we also tested another approach that formulates the age prediction task as a classification problem, rather than a regression one. The idea was to investigate whether the integration of a classifier into the age prediction algorithm could help diminish the intrinsic issue of “regression towards the mean” that characterises regression models (Liang et al., 2019), i.e., the tendency of overestimating the predicted age of younger subjects and underestimating it in older subjects. Moreover, the availability of classification probability estimates—produced as output by the classifier—could help to weigh the importance of the LR prediction when generating the final predicted age.

The whole available age range (i.e., between 14 and 70 weeks) was split into 40 bins and each subject was assigned to its relative “age bin”, which describes the age to be predicted by the model. Of all the 40 evenly spaced bins, 32 were empty (i.e. they did not correspond to any of the ages of the training subjects), while the remaining 8 age bins were represented by three or more subjects. Thus, these 8 bins could be used as classes to train a multinomial LR classifier. Moreover, since the data set is affected by class imbalance, different weights were associated to each class by assigning values that are inversely proportional to the class frequencies.

Once the LR-based age predictions were obtained, their probability estimates were used to calculate a weighted average between the LR and the GPR predictions. If *Ŷ _GPR_* is the age predicted by the GPR model on an input subject, *Ŷ*_*LR*_ is the most likely age class predicted by the LR model on the same subject and *p*_*LR*_ is the corresponding probability estimate, then the final predicted age *Ŷ* was simply calculated as:

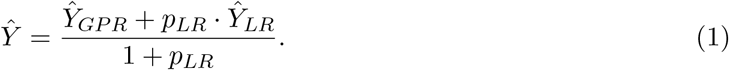

The performance of the implemented age prediction model was evaluated by computing both the MAE and the linear correlation between chronological and predicted ages on the validation sets.

Furthermore, the trained LR classifier also provides the weights for each input feature in the decision function. Therefore, we used the average weights for all features to investigate feature importance for age classification. We did so by, first, multiplying the weights by the standard deviation of the corresponding feature in the input data. Finally, we divided each of these values by their total sum (calculated across all features), in order to obtain a more interpretable “relative” importance score with respect to all available input features.

### 2.6 Testing on the ageing cohort

Both the GPR and LR models were re-trained one last time including all 31 training subjects together. The proposed pipeline was then employed for inference on both the control and EEDR rats of the ageing cohort. MAE and correlation between chronological and predicted ages were computed for this data set, and differences between correlations computed on the same observations were tested with the method proposed by Steiger for comparing dependent correlations (Steiger, 1980). Moreover, for each subject and at each time point, the BrainAGE score was computed as the difference between predicted and chronological age.

A linear mixed effect model was fitted by setting the BrainAGE score as the dependent variable, with chronological age, lifestyle (controls vs. EEDR) and their interaction as fixed effects. Moreover, the ID of each subject was set as a random effect to account for repeated measures. This analysis was carried out in order to test for group differences across time between controls and EEDR rats.

Finally, Cox regression was employed to investigate the effects of lifestyle and BrainAGE scores at session 1 (corresponding to ages between 12 and 14 weeks) and session 2 (49 to 51 weeks) on total survival. This was done by fitting five different regression models, using, respectively, the following five sets of independent variables: (1) both lifestyle group and BrainAGE score at session 1; (2) both lifestyle group and BrainAGE score at session 2; (3) only lifestyle group; (4) only BrainAGE score at session 1; (5) only BrainAGE score at session 2. The other two available time points (session 3 and 4) were not included in the survival analysis, since several of the subjects were discarded from the study after the second session.

## 3 Results

### 3.1 Performance of the age prediction model on the training cohort

In the first part of the study, the age prediction model was designed and evaluated within the training cohort using leave-one-out cross-validation. This allowed us to investigate which inputs and model configuration led to the best performance, as well as which image features were important for age prediction.

#### 3.1.1 Input and performance of the GPR model

The age prediction accuracy was first evaluated by testing different combinations of model inputs. As already mentioned in Section 2.5.1, our final choice was to feed the model with all three TPMs, modulated using the 11 months reference template only. This configuration led to the lowest MAE in the age predictions, which was equal to 5.69 weeks (see Figure 1a). All other tested inputs—including the concatenation of the TPMs modulated after registration to all four available templates—showed comparable or slightly higher MAEs (see Supplementary Material).

**Figure 1:**
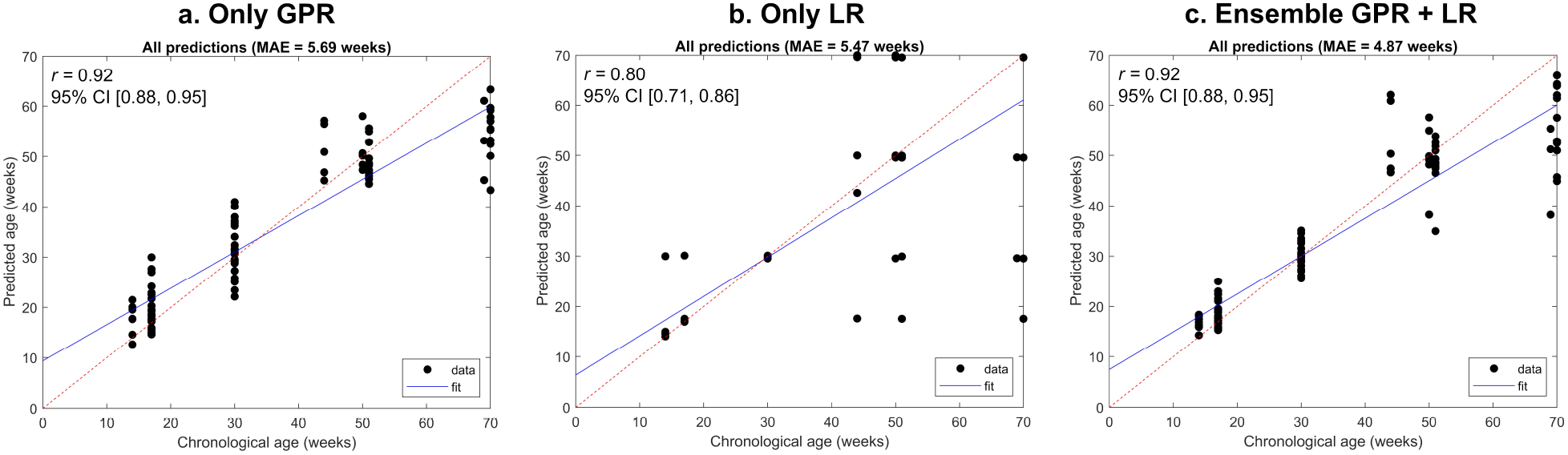
Comparison of the performance of the three analysed models (a: GPR only, b: LR classifier only, c: ensemble of GPR and LR) for age prediction. In each plot, the age predicted from each input sample is represented on the *y* axis, against the actual chronological age on the *x* axis. The correlation coefficient between predicted and chronological age, together with its 95% confidence interval, is also presented for each model. All predictions (estimated with leave-one-out cross-validation) are represented with black dots, while their fitted line is shown in blue. The red dashed line corresponds to what would be the “perfect” prediction (i.e. predicted age equal to chronological age).

We then calculated the linear correlation coefficient between predicted and chronological ages for the chosen model, which was equal to 0.92 (95% confidence interval = [0.88, 0.95]).

#### 3.1.2 Comparison of different models

Once the model inputs had been set, we compared the performance of the GPR model with that of both the LR classifier and the proposed ensemble of GPR and LR. As shown in Figure 1, the three models performed differently.

The LR classifier alone led to a slightly lower MAE compared to GPR (5.47 vs. 5.69 weeks), but also to a worse correlation coefficient between predicted and chronological ages, i.e. 0.80 (95% confidence interval = [0.71, 0.86]) vs. 0.92 (95% confidence interval = [0.88, 0.95]). However, this difference in correlation coefficients did not turn out to be statistically significant (*p* = 0.08). From the plot in Figure 1b, it can be noticed that the main improvements in prediction accuracy were obtained at younger ages (especially at session 2, where predictions were perfectly accurate). Moreover, the fitted line in the plot shows a slight reduction in the tendency of making predictions “towards the mean”. However, the LR prediction errors were far greater for the later scan sessions compared to GPR.

The best performance was observed by weighing together the predictions from GPR and LR, as shown in Figure 1c. The lowest MAE was indeed observed (equal to 4.87 weeks), while still maintaining a correlation of 0.92 (95% confidence interval = [0.88, 0.95]) between predicted and chronological ages. For this reason, we decided to use this ensemble model for future testing on the ageing cohort. The individual ageing trajectories obtained using such a model in the training cohort are shown in Figure 2.

**Figure 2:**
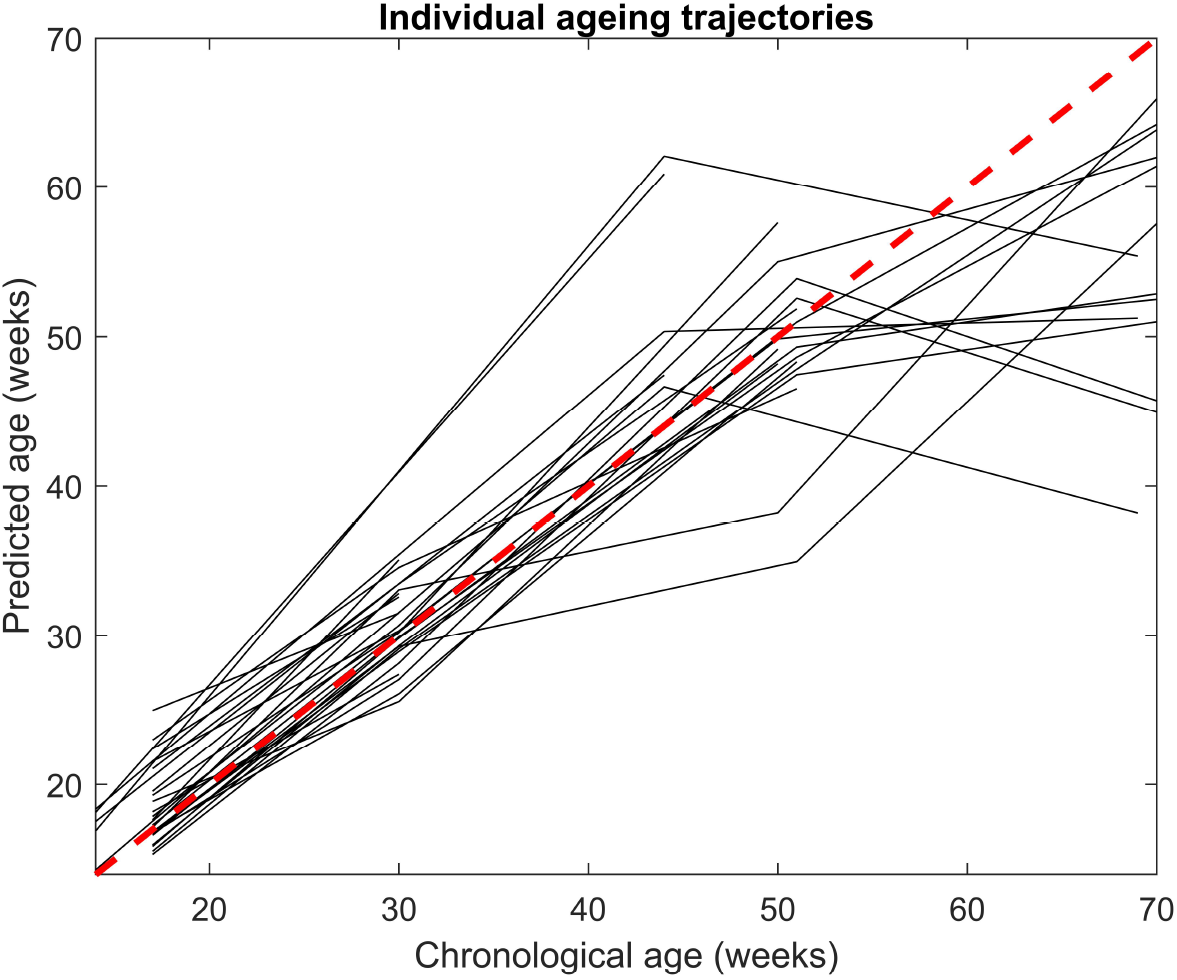
Age trajectories (in black) estimated for all 31 subjects in the training cohort using the proposed ensemble model of GPR and LR. The red dashed line corresponds to what the trajectory would be in case of predictions that are perfectly matching with the corresponding chronological ages.

#### 3.1.3 Feature importance analysis

We investigated the most important input features for age classification by ranking the relative importance scores from the highest to the lowest. Each of these analysed features represents one of the 77 PCs extracted through principal component analysis. The highest scores were found for feature 2 (normalised importance score of 10%) and feature 1 (score of 7.8%). They were followed by feature 5 (4.2%) and feature 4 (3.7%). All other features had scores equal to or lower than 2.8%. After a first exponential drop in feature importance across the first 10 features, all other importance scores decreased with an approximately linear decay until reaching the lowest score of all (equal to 0.2%).

We transformed the model’s inputs back into the original image space and visualised the distribution of the features in the modulated tissue maps. Figure 3 shows a representation of the main variations in the modulated TPMs for the two most important components for age classification (PC 2 and 1). From the figure, it can be noticed that a few areas are affected by more changes than others (e.g. parts of the cerebellum, the amygdala and the hippocampus for feature 2, or the brain stem for feature 1). However, in general, PC 2 and 1 seem to represent spread variations of various magnitude throughout the brain, rather than in very specific regions of interest (ROIs). Similar unspecific patterns could also be observed for other features, which followed feature 2 and 1 in the importance score ranking.

**Figure 3:**
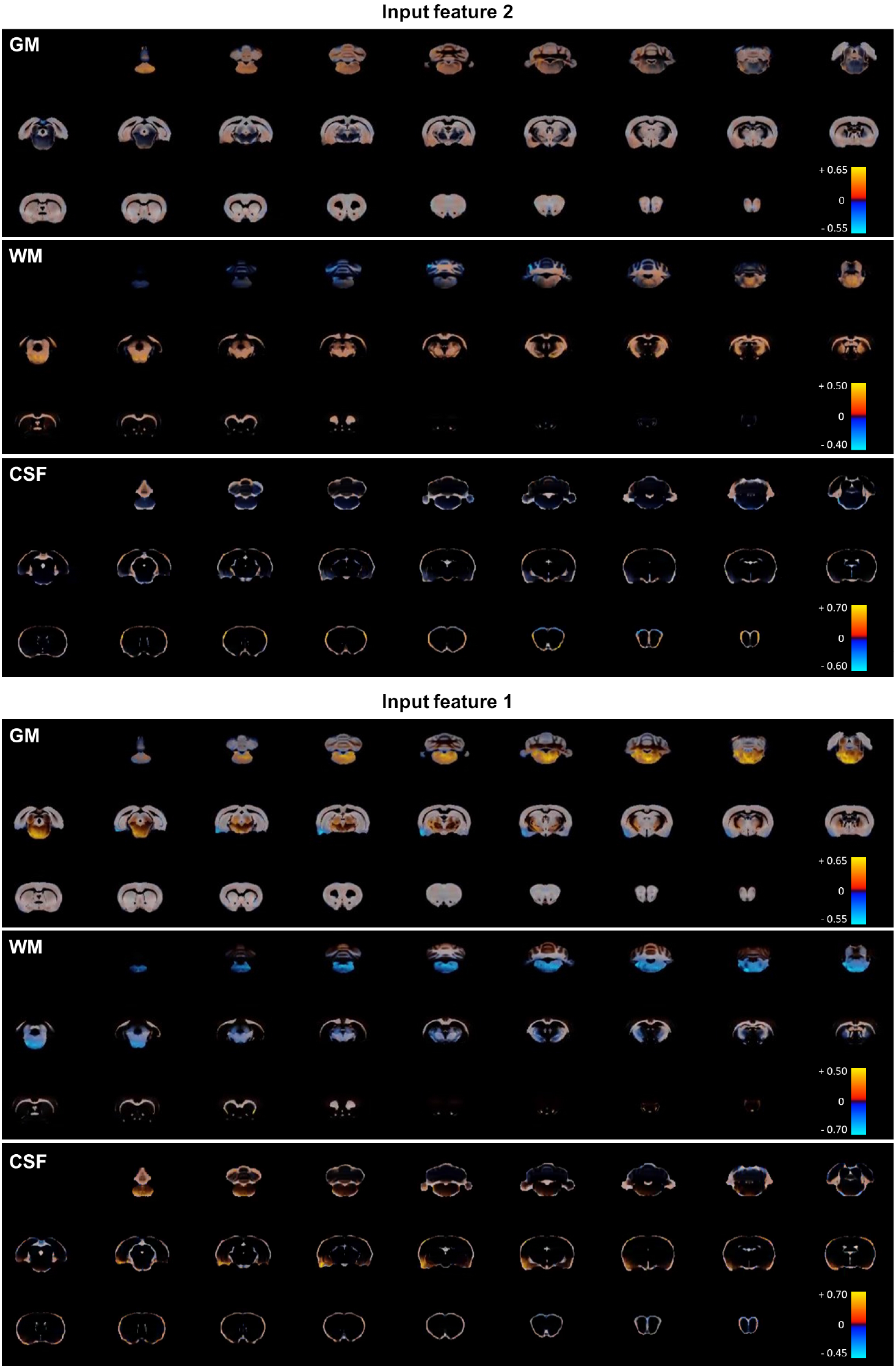
The main GM, WM and CSF variations represented by feature 2 (top) and feature 1 (bottom) are shown. For each PC (1 or 2) and each type of modulated TPM (GM, WM or CSF), the PC’s standard deviation multiplied by 2 (corresponding to one of the two extremes of the PC’s distribution) is overlaid on the relative mean TPM. The opacity of the double standard deviations is modulated by intensity, making more visible the regions that differ the most from the mean. Higher and lower values with respect to the mean are colour-coded using, respectively, shades of red and blue.

### 3.2 Effect of intervention on the frailty of the ageing cohort

In order to confirm that EEDR intervention was successful in improving the general health status of the rats in the ageing cohort, we measured their frailty index (FI). The distributions of the scores between the groups and across time are presented in Figure 4. The EEDR group at session 3 had a mean FI 1.5 ± 0.9, while the mean for the control group at the same session was 2.7 ± 1.3. At session 4, the mean FI of EEDR rats was 1.2 ± 0.8, while the control rats’ FI was 2.4 ± 1.4. A linear mixed-effects model was used to evaluate the difference in frailty between the two groups at both time points. The main effect of scan session was not significant (F(1,30) = 0.7149, *p* = 0.4045), and there was no significant interaction between the main terms (F(1,30) = 0.06706, *p* = 0.7974). There was, however, a significant main effect of the group (F(1,37) = 18.15, *p* = 0.0001). Post-hoc comparisons show that the controls had a higher frailty score at both session 3 (95% confidence interval of difference = [0.4829, 2.057], *p* = 0.0009) and session 4 (95% confidence interval of difference = [0.4873, 2.253], *p* = 0.0014) relative to the EEDR rats. These results show that, in later life, the control rats were in worse overall physical health than the EEDR rats but that there were some overlaps between the groups.

**Figure 4:**
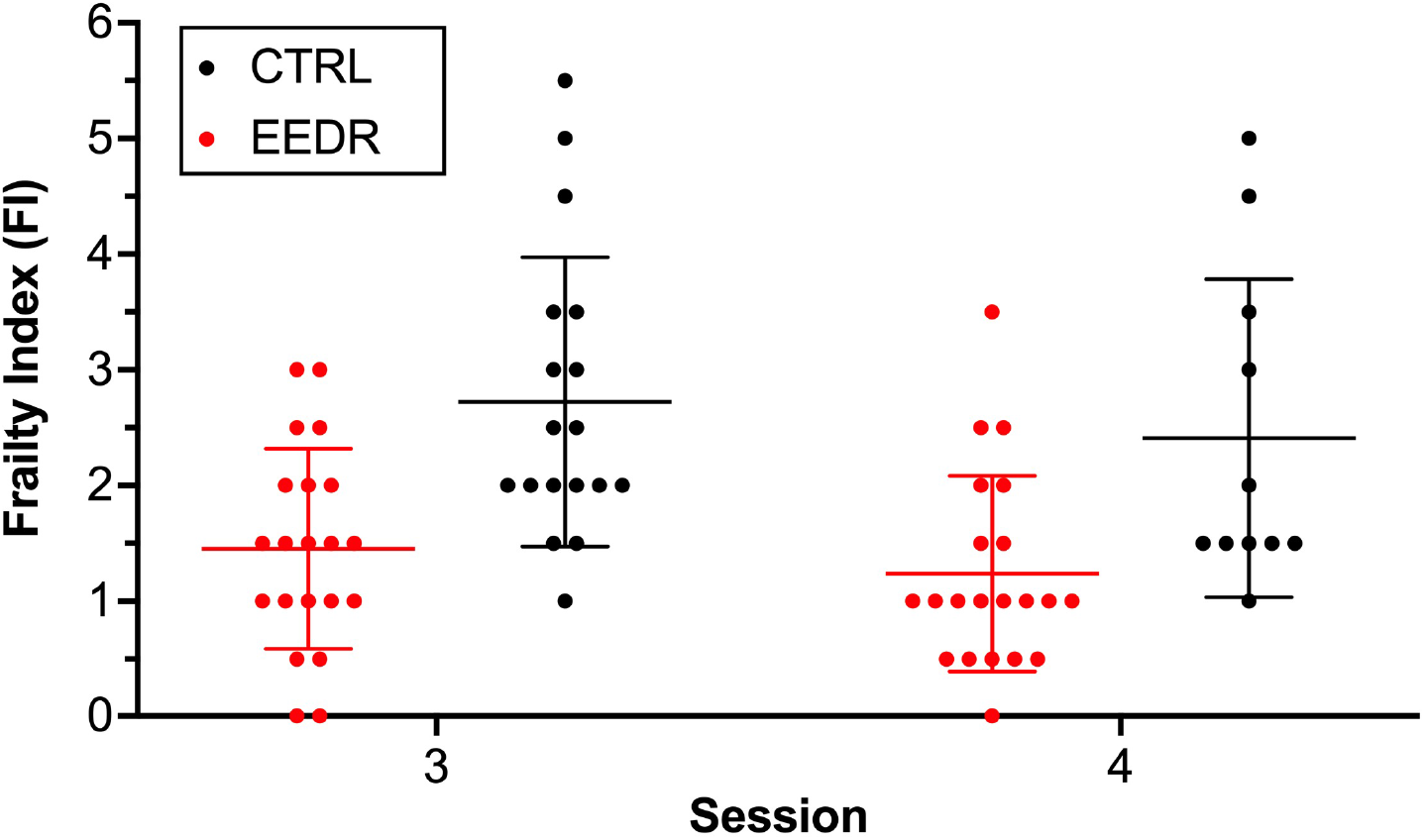
The frailty index (FI) for subjects in the ageing cohort immediately before scanning session 3 (50 ± 1 weeks) and 4 (73 ± 1 weeks). The error bars denote the mean ± the standard deviation. The individual scores are plotted as a black or a red dot depending on whether the input subject belongs to the control or the EEDR group, respectively. There was a significant main effect of group (F(1,37) = 18.15, *p* = 0.0001), but not of session (F(1,30) = 0.7149, *p* = 0.4045) or interaction (F(1,30) = 0.06706, *p* = 0.7974), according to the mixed effects model.

### 3.3 Age prediction on the ageing cohort

Once the age prediction model was trained, it was employed to predict the ages of the rats from the ageing cohort. Figure 5 shows the age predictions obtained on that cohort, distinguishing control subjects from the EEDR ones.

**Figure 5:**
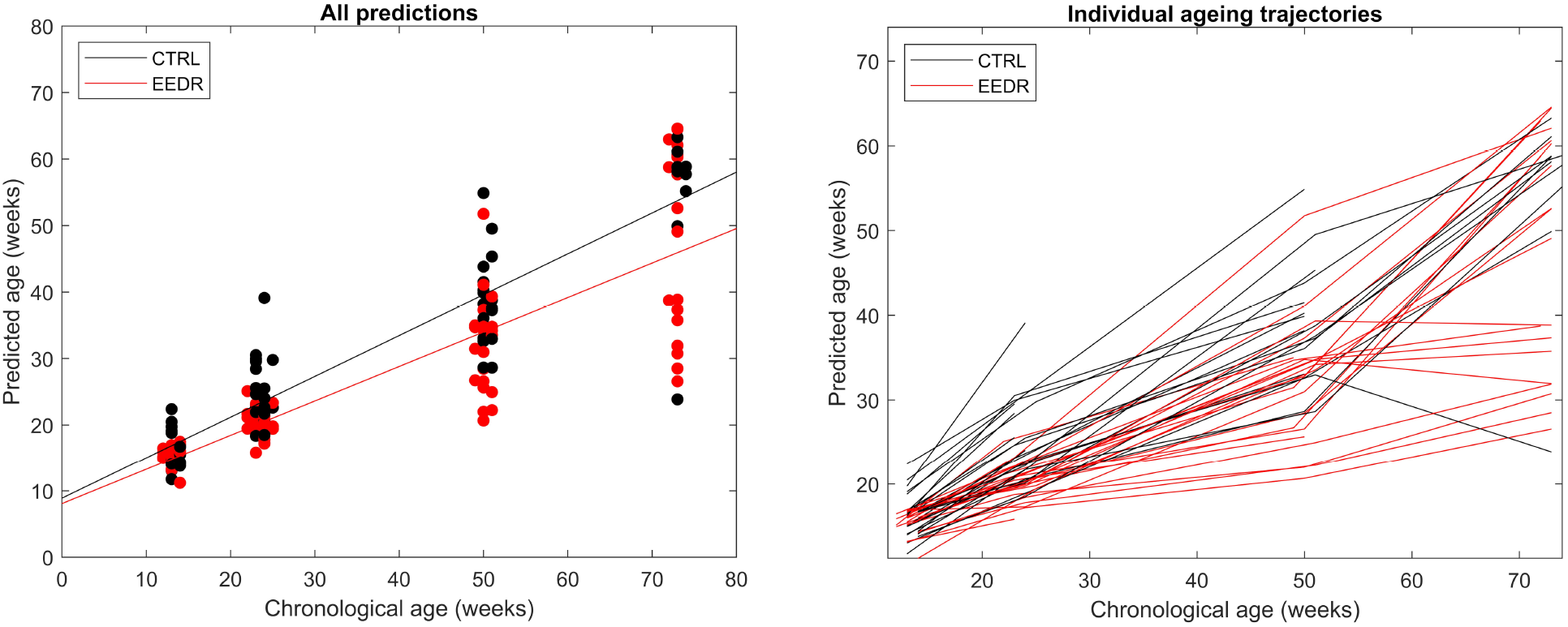
Age prediction results obtained on the test subjects from the ageing cohort. On the left, each prediction is plotted as a black or a red dot depending on whether the input subject belongs to the control or the EEDR group, respectively. The black and red lines show the linear fitting for each of the two groups, having slopes of 0.61 (controls) and 0.52 (EEDR). On the right, the estimated ageing trajectories for all subjects are plotted using the same colour-code in order to distinguish the two groups.

A decrease in the prediction accuracy was observed by testing the model on the new cohort. The MAE was 9.89 weeks when considering all subjects together, while it was 7.47 weeks for the control group alone, and 11.88 for the EEDR group only.

The correlation coefficient between chronological and predicted age also decreased to 0.86 (95% confidence interval = [0.81, 0.89]) for all subjects together. This result is mainly influenced by the EEDR samples, which alone showed a correlation coefficient of 0.85 (95% confidence interval = [0.78, 0.90]), against 0.91 (95% confidence interval = [0.85, 0.94]) for the control group.

#### 3.3.1 Group differences in BrainAGE score

For all subjects and all samples in the ageing cohort, we computed the respective BrainAGE score. A linear mixed-effects model was then fitted in order to investigate group differences across time.

Statistically significant effects were observed for chronological age alone (*b* = −0.48, 95% confidence interval = [−0.54, −0.43], SE = 0.03, *t* = −17.39, *p* < 0.001) and the interaction between lifestyle (i.e. EEDR or control) and chronological age (*b* = 0.11, 95% confidence interval = [0.02, 0.20], SE = 0.04, *t* = 2.47, *p* = 0.015). On the other hand, lifestyle alone did not show any significant effect on the BrainAGE score (*p* = 0.777).

#### 3.3.2 Survival analysis

Cox regression models were fitted to perform survival prediction using lifestyle information and BrainAGE scores at sessions 1 and 2. The proportional hazard assumption was met by the model, supporting the use of Cox regression.

First, we fitted three models with just one independent variable each. No significance was found for BrainAGE (*p* = 0.102) or for lifestyle (*p* = 0.0503) at session 1, i.e. before EEDR intervention began. On the other hand, BrainAGE score at session 2 showed a significant effect on survival prediction (*p* = 0.03). In particular, lower BrainAGE scores at session 2 correlated with longer survival (see Figure 6).

**Figure 6:**
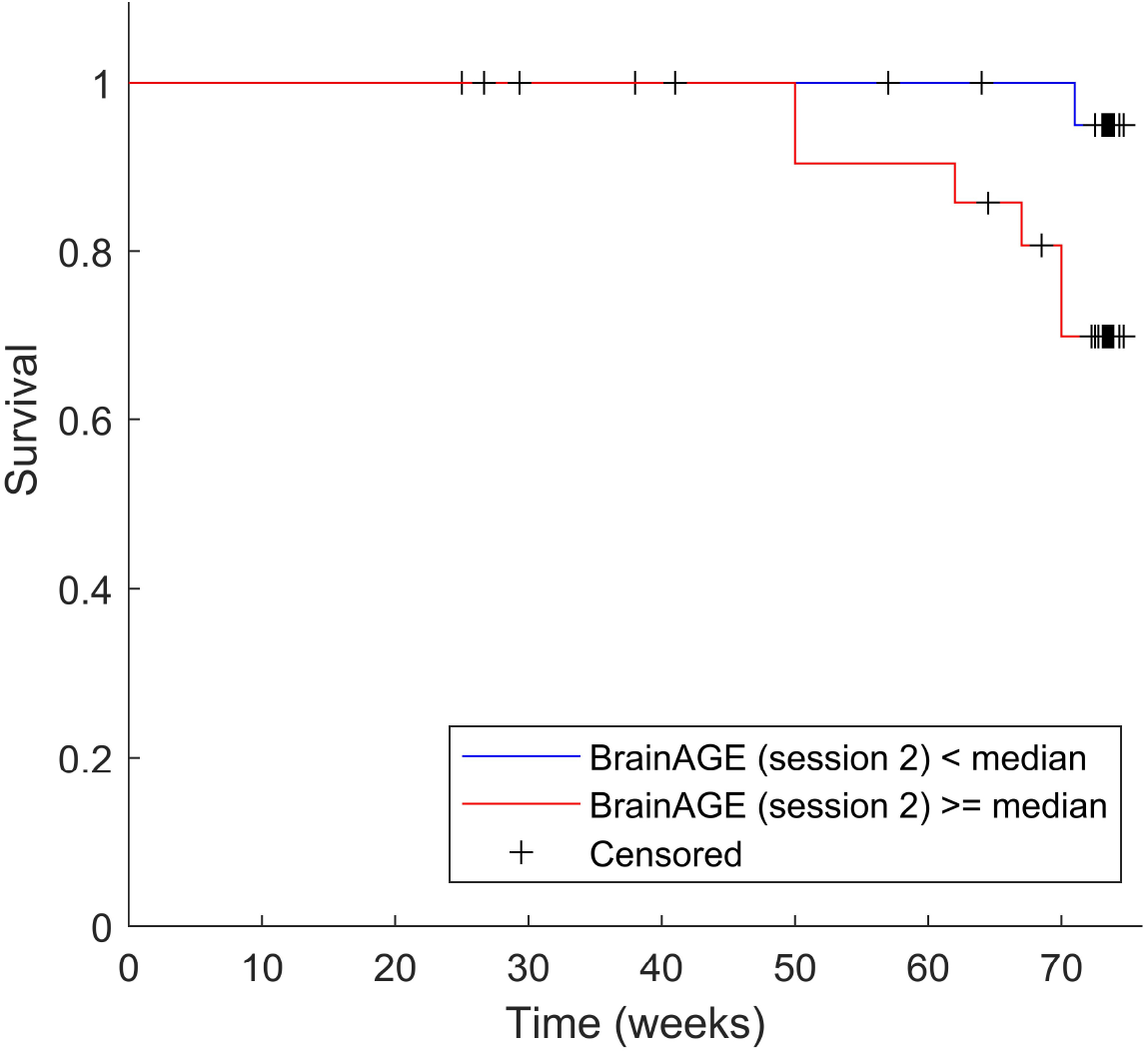
Estimated survival for the subjects included in the ageing cohort. Subjects with a BrainAGE score at session 2 (i.e. between 22 and 25 weeks old) that is lower than the median are shown to have higher chance of survival until the end of the observation period of the present study.

Two additional regression models were fitted using two independent variables (lifestyle and BrainAGE score at session 1 together, as well as lifestyle and BrainAGE score at session 2). How-ever, they did not add any relevant information to the survival analysis, since no variable showed a statistically significant result.

## 4 Discussion

In this work, we propose and validate a novel MRI-based pipeline for brain age prediction on rats. The core of the experiment was the controlled modulation of the ageing process by an active and healthy lifestyle. We verified the premise that EEDR intervention significantly improves the rats’ overall health by demonstrating a significantly lower FI at two latter time points (approximately 50 and 73 weeks of age). The results of other parameters collected during this longitudinal imaging and behavioural study will be presented elsewhere (manuscripts in preparation).

### 4.1 Model performance on the training cohort

The performance of our proposed model was first investigated by training and validating it on a training cohort of rats, which produced accurate prediction results.

We believe that the choice of GPR as a baseline model was suitable for our preclinical validation, since it already showed accurate results in multiple human studies (Cole et al., 2015, 2018, 2017b). Furthermore, previous brain age studies have not found significant differences in performance between GPR and relevance vector regression—which is the model used on rats by Franke et al. (2016)—when trained on the same data type (Aycheh et al., 2018; Baecker et al., 2021). Thus, we considered it reasonable to only test GPR as a baseline for the present study. Finally, given the limited size of our training set, we did not test deep learning-based approaches, despite their successful outcome in previous human studies (Cole et al., 2017b; Jónsson et al., 2019; Bashyam et al., 2020).

Compared to the previous study by Franke et al. (2016) on rat brain age prediction, our model showed a very similar accuracy using leave-one-out cross-validation. By integrating an LR classifier into the pipeline, we obtained a MAE of 4.87 weeks (corresponding to approximately 34 days and 8.7% of the total available age range), while Franke et al. (2016) reported a MAE of 49 days (equating to a mean error of 6% with respect to their used age range). On the other hand, they presented a linear correlation coefficient of 0.95 between chronological and predicted age, while we obtained a coefficient of 0.92. This small loss in performance may partially stem from the difference in size and heterogeneity of the data sets: Franke et al. (2016) implemented the model using a total of 273 scans from 24 subjects, as opposed to our available 89 scans from 31 subjects. However, despite these differences in study design, the results of our work are rather consistent with Franke et al. (2016), further supporting the potential of applying brain age prediction models across laboratories.

#### 4.1.1 Integration of a classifier into the age prediction workflow

Compared to previously-proposed brain age prediction models (in both humans and animals), the main methodological novelty presented in this work consists in the integration of a classifier into the age prediction task. By dividing the age range into separate classes, a classifier can then assign each input to its most appropriate age class, thus avoiding the known bias of regression approaches to make predictions that tend to be closer to the mean. As presented in Section 3.1.2 and Figure 1, the classifier alone does not perform better than the traditional regression approach. However, it is possible to observe that certain predictions are particularly accurate, especially at younger ages, and such predictions are also more likely to present higher probability estimates. Accordingly, our results showed that performing a weighted average between GPR and LR predictions—by weighing the LR estimations with their relative probability estimates—can decrease the age prediction error in the training set, while also maintaining a satisfactory correlation between predicted and chronological age.

#### 4.1.2 Difference in prediction accuracy between younger and older ages

As mentioned above and represented in Figures 1 and 2, the age predictions are particularly accurate at younger ages, while at the later two sessions the prediction error tends to increase. However, this result was expected for two main reasons.

First of all, as presented in Section 2.1, the earlier sessions had a higher number of image samples compared to the later ones. Therefore, it is reasonable to expect an influence of such an imbalanced distribution in the final accuracy across different ages.

Furthermore, we believe that the discrepancy in performance may also reflect actual anatomical differences between subjects that become more and more accentuated with time. While the brain morphology of all subjects may be relatively similar in the earliest months of life, their ageing process in later life can be significantly affected by various environmental or genetic factors (Cole et al., 2018). This may lead to a greater variability in brain ageing trajectories at older ages, making it difficult to establish whether a larger MAE in age prediction is actually a sign of poor model performance, rather than simply reflecting individual differences in the ageing process. Other data from this study (MacNicol et al., 2019) suggest the latter to be the case. Similar patterns were also observed in the previous study by Franke et al. (2016), where a higher variance was observed at older ages in the individual ageing trajectories of 23 untreated rats.

As can be observed from Figure 2, the steepness of the ageing trajectories tends to decrease between the third and the fourth scan sessions in the majority of subjects. We believe that to be the evidence of non-linearity in the brain ageing process, which is in accordance with previous longitudinal studies that reported non-linear trajectories of both structural and functional brain data (Raz et al., 2005; Pfefferbaum et al., 2013; Fjell et al., 2013; Vinke et al., 2018). On the other hand, it should also be taken into consideration that the animals with a more rapid ageing process were more likely to be lost or excluded at follow-up, due to either dying, obesity or health problems.

#### 4.1.3 Feature importance for age classification

The analysis of the LR model’s coefficients allowed us to investigate which of the input PCs are more important for age classification. We show that it was not possible to identify specific focal areas that strongly correlate with age prediction, but rather widespread variations were found across the whole brain.

Diffuse brain changes related to ageing have previously been observed in human studies. For example, Good et al. (2001), using voxel-based morphometry, reported widespread GM changes during ageing, involving cortical regions, deep GM structures and the cerebellum. On the other hand, they did not find relevant changes in WM volume, which we show to be affected by widespread changes in the present study. Another study by Walhovd et al. (2005) investigated the effects of ageing by comparing the volumes of different regions of interest (of GM, WM and CSF) and also identified significant differences in almost all the analysed volumes. Moreover, these differences were found to be heterogeneous across various ROIs that also match the findings of our study (see Figure 3): some volumes showed a linear relationship with age (e.g. amygdala, thalamus and cortex), but others a curvilinear relationship (e.g. hippocampus, brainstem and cerebellum). As we point out (Turkheimer et al., 2021), these complex patterns observed in longitudinal structural data may stem from nonlinear interactions that are highly influenced by various environmental, metabolic and immune factors, which can vary across time. This concept further supports the idea that the whole brain—rather than only specific regions—is affected by nonlinear changes that contribute to the ageing process, and that such changes are rather heterogeneous between and within subjects through time.

### 4.2 Effect of the EEDR lifestyle on BrainAGE

Once the age prediction model had been trained, it was tested on an independent cohort consisting of rats with one of two lifestyles: controls (which had no intervention and were comparable to the rats in the training cohort) and EEDR. To our knowledge, this is the first study to perform an active lifestyle intervention on rats with the aim of analysing its effect on brain age prediction. From the age estimations obtained on this cohort (Figure 5), two major conclusions can be drawn.

First of all, a general bias is present in the age estimations, which tend to be lower than what was observed in the training cohort (see Figure 1), especially at older ages. This loss in accuracy was also reflected in the MAE and the correlations between predicted and chronological ages (see Section 3.3). However, we believe that a loss in performance is to be expected when testing machine learning-based models on new unseen cohorts that inevitably differ from the training set, especially when the size of the training data set is rather limited. The decreased accuracy at older ages still reflects the patterns that were observed on the training cohort, as previously discussed (Section 4.1.2). In addition, a higher MAE and a lower correlation coefficient between predicted and chronological ages were found on EEDR rats compared to controls. This difference between groups supports the hypothesis that the controls were more comparable to the subjects in the training data set despite belonging to a separate cohort.

The second main outcome observed from our results is that there is a difference between the two lifestyle groups in terms of their average ageing trajectories, and that this difference increases over time. This was confirmed by the linear mixed-effects model, in which not only chronological age, but also the interaction between chronological age and lifestyle, had a significant effect on the BrainAGE score. As there was no significance for the simple lifestyle group factor, this means that the two groups did not have significantly different BrainAGE scores at young chronological age but aged at different average speeds depending on their lifestyle. This result is of high relevance, since it shows that a healthier lifestyle—in this case characterised by a better diet and an enriched environment—may potentially slow down or delay the ageing process. Previous studies on both rodents and humans have already reported a strong association between ageing outcome and dietary habits (Mattson, 2010; Martin et al., 2006; Maioli et al., 2012; Soininen et al., 2021), physical exercise (Hötting and Röder, 2013), environmental enrichment (Speisman et al., 2013), as well as multidomain lifestyle interventions (Ngandu et al., 2015). Moreover, our results are in accordance with a recent study (Bittner et al., 2021) that, to the best of our knowledge, constitutes the most thorough attempt at investigating the relationship between BrainAGE score and multivariate lifestyle behaviours in humans (i.e., alcohol consumption, smoking, social integration and physical activity). They indeed showed that lifestyle habits do affect brain age estimations, with smoking and lower physical activity contributing the most to this association. The consistency between the findings from preclinical and human studies is of fundamental importance for strengthening the case for using BrainAGE as a valid ageing biomarker. Moreover, it is possible to better perform active interventions, such as dietary restrictions, and control for specific lifestyle factors within a preclinical framework compared to human studies (at least for a life-long observation period).

According to our results, though, it is also important to point out that the BrainAGE score alone does not provide relevant information on how healthy the ageing of an individual is, since we demonstrated that its value is strongly affected by the chronological age. Similar results have also been observed in previous human studies and are largely related to the problem of age regression towards the mean (Liang et al., 2019; Cole et al., 2017b; Pardoe and Kuzniecky, 2018). However, if chronological age is also taken into account, it may be possible to draw more informed conclusions from the BrainAGE score. The pairing of chronological age and BrainAGE information could indeed represent an indicator of the brain ageing process of individuals, by comparing their BrainAGE scores with what is expected from a control group at the same chronological age.

### 4.3 Survival analysis

The potential of using the BrainAGE score as a biomarker for healthy ageing was further strengthened by the results of the survival analysis (Section 3.3.2). BrainAGE at session 2 (i.e. before any subjects were excluded from the study because of death or other reasons) was shown to have a significant effect on survival. In particular, subjects presenting a lower BrainAGE score between 22 and 25 weeks of age were found to be more likely to survive longer than those with higher scores. This finding is in agreement with a previous human study by Cole et al. (2018) that reported a significant correlation between BrainAGE and mortality before the age of 80, an association that was also shown to be independent from other possible influences (e.g. education, social class, or the presence of age-associated illness).

On the other hand, we found no corresponding significant effect on survival of BrainAGE score at session 1 (before EEDR intervention began, when the variation between individuals is less pronounced) or of lifestyle group. As discussed in the previous section, the EEDR lifestyle was shown to have a significant effect on the BrainAGE score. However, lifestyle information alone does not appear to be sufficient for predicting survival. This result highlights the importance of using the proposed MRI-based predictions for investigating mortality.

### 4.4 Limitations and future work

The present study is affected by some limitations that we aim to overcome in the future. The first drawback consists in the limited data set, especially when it comes to the training cohort. This is a common problem of animal studies, and we believe that the accuracy of our proposed age prediction model might improve by increasing the size of the training data set. Moreover, our current training cohort includes only male rats, while most human brain age prediction studies include both male and female subjects and account for sex as a covariate in the data analyses (Cole et al., 2015; Pardoe et al., 2017; Cole et al., 2017c). Thus, in the future, we would like to use more image samples during training, which should include both male and female rats. We would also aim to scan such rats more often (rather than just from four sessions) and over a longer observation period than the one used here. This could allow us to gain a better insight on brain age prediction throughout the lifespan. Furthermore, an increase in training samples would also strengthen the potential of successfully using our method on data acquired from different sites and at different ages, without having to re-train the model. Finally, an increase in data samples in the test cohort too could allow us to more thoroughly investigate the value of BrainAGE as a biomarker. For example, a longer study observation period and the occurrence of more death events may strengthen the power of the survival analysis.

In the present work, we used the LR model coefficients to get a better understanding of the brain regions that influence age prediction the most. However, when it comes to GPR, the use of a linear covariance function could not allow us to extract measures that could directly relate with feature importance for the regression model. In the future, we aim to investigate the effect of using alternative kernels with automatic relevance determination. This approach consists in including a length-scale parameter for each input feature within the covariance function, and these parameters could later be analysed to determine feature importance for prediction (Caywood et al., 2017). Moreover, we would also like to explore the use of ROI-based measurements (e.g. regional volumes obtained through atlas-based segmentation) for age prediction. This could allow us not only to investigate whether ROI information can improve prediction accuracy, but also to study the influence of each ROI separately on age prediction.

Finally, we would ideally test the proposed age prediction model on rodent models of abnormal, pathological ageing. This would enable us to identify if and how early the BrainAGE score— computed using the present strategy—can reveal anomalies in the rodents’ ageing process, similarly to what has already been performed in previous studies on humans undergoing neurodegeneration. This would allow us to gain a better insight in the potential of BrainAGE as an ageing biomarker within a controlled preclinical framework, and to utilise this in testing novel and experimental treatments for neuropsychiatric disorders.

## 5 Conclusion

We present a new algorithm for rodent brain age prediction, based on neuroimaging and machine learning. Using the proposed method, which integrates regression and classification, we achieved high prediction accuracy on a training cohort of control rats, supporting the potential of using structural MRI data for extracting accurate information on brain age. Furthermore, to the best of our knowledge, this is the first preclinical work to test such a prediction model on a new separate cohort of animals. We investigated predicted age and BrainAGE scores on two additional groups of rats: controls and subjects that underwent EEDR. Our results indicate that EEDR significantly affects the ageing trajectories of the analysed rats by slowing down their ageing process. Moreover, the BrainAGE score at approximately 5 months of age was found to have a significant effect on survival. These findings are in agreement with previous studies on humans and support the potential of using MRI-based brain age prediction models as a biomarker of healthy ageing.

## Acknowledgements

The authors would like to thank Dr Francesca Biondo for her expertise and support during data preprocessing, as well as Sebastian Popescu for his assistance and initial inspiration for the implementation of the Python-based GPR model.

## Author contributions

**Irene Brusini**: Conceptualisation, Methodology, Software, Validation, Formal analysis, Writing - Original Draft, Visualisation. **Eilidh MacNicol**: Conceptualisation, Methodology, Investigation, Resources, Data Curation, Writing - Review & Editing. **Eugene Kim**: Resources, Data Curation, Writing - Review & Editing. **Ö rjan Smedby**: Formal analysis, Writing - Review & Editing. **Chunliang Wang**: Methodology, Writing - Review & Editing. **Eric Westman**: Conceptualisation, Writing - Review & Editing. **Mattia Veronese**: Conceptualisation, Methodology, Writing - Review & Editing, Supervision, Funding acquisition. **Federico Turkheimer**: Conceptualisation, Methodology, Writing - Review & Editing, Supervision, Funding acquisition. **Diana Cash**: Conceptualisation, Writing - Review & Editing, Supervision, Project administration, Funding acquisition.

## Data and code availability statement

The presented Python-based age prediction pipeline can be found at https://github.com/ibrusini/RAGE. The resources generated for this study (metadata, trained models, age estimations) can be found in a separate Open Science Framework repository (DOI: 10.17605/OSF.IO/6WD7T).

## References

Ashburner, J., Friston, K.J., 2000. Voxel-based morphometry—the methods. NeuroImage 11, 805–821.

Avants, B.B., Tustison, N.J., Wu, J., Cook, P.A., Gee, J.C., 2011. An open source multivariate framework for n-tissue segmentation with evaluation on public data. Neuroinformatics 9, 381–400.

Aycheh, H.M., Seong, J.K., Shin, J.H., Na, D.L., Kang, B., Seo, S.W., Sohn, K.A., 2018. Biological brain age prediction using cortical thickness data: A large scale cohort study. Frontiers in Aging Neuroscience 10. doi:10.3389/fnagi.2018.00252.

Baecker, L., Dafflon, J., Da Costa, P.F., Garcia Dias, R., Vieira, S., Scarpazza, C., Calhoun, V.D., Sato, J.R., Mechelli, A., Lopez Pinaya, W., 2021. Brain age prediction: A comparison between machine learning models using region-and voxel-based morphometric data. Human Brain Mapping.

Bashyam, V.M., Erus, G., Doshi, J., Habes, M., Nasralah, I., Truelove-Hill, M., Srinivasan, D., Mamourian, L., Pomponio, R., Fan, Y., 2020. Mri signatures of brain age and disease over the lifespan based on a deep brain network and 14 468 individuals worldwide. Brain 143, 2312–2324.

Bittner, N., Jockwitz, C., Franke, K., Gaser, C., Moebus, S., Bayen, U.J., Amunts, K., Caspers, S., 2021. When your brain looks older than expected: combined lifestyle risk and brainage. Brain Structure and Function doi:10.1007/s00429-020-02184-6.

Burns, A., Iliffe, S., 2009. Alzheimer’s disease. Bmj 338, b158. doi:10.1136/bmj.b158.

Caywood, M.S., Roberts, D.M., Colombe, J.B., Greenwald, H.S., Weiland, M.Z., 2017. Gaussian process regression for predictive but interpretable machine learning models: An example of predicting mental workload across tasks. Frontiers in Human Neuroscience 10. doi:10.3389/fnhum.2016.00647.

Cole, J.H., Annus, T., Wilson, L.R., Remtulla, R., Hong, Y.T., Fryer, T.D., Acosta-Cabronero, J., Cardenas-Blanco, A., Smith, R., Menon, D.K., 2017a. Brain-predicted age in down syndrome is associated with beta amyloid deposition and cognitive decline. Neurobiology of aging 56, 41–49.

Cole, J.H., Franke, K., 2017. Predicting age using neuroimaging: innovative brain ageing biomarkers. Trends in Neurosciences 40, 681–690.

Cole, J.H., Leech, R., Sharp, D.J., Initiative, A.D.N., 2015. Prediction of brain age suggests accelerated atrophy after traumatic brain injury. Annals of neurology 77, 571–581.

Cole, J.H., Poudel, R.P., Tsagkrasoulis, D., Caan, M.W., Steves, C., Spector, T.D., Montana, G., 2017b. Predicting brain age with deep learning from raw imaging data results in a reliable and heritable biomarker. NeuroImage 163, 115–124.

Cole, J.H., Ritchie, S.J., Bastin, M.E., Hernández, M.V., Maniega, S.M., Royle, N., Corley, J., Pattie, A., Harris, S.E., Zhang, Q., 2018. Brain age predicts mortality. Molecular psychiatry 23, 1385–1392.

Cole, J.H., Underwood, J., Caan, M.W., De Francesco, D., van Zoest, R.A., Leech, R., Wit, F.W., Portegies, P., Geurtsen, G.J., Schmand, B.A., 2017c. Increased brain-predicted aging in treated hiv disease. Neurology 88, 1349–1357.

Denver, P., McClean, P.L., 2018. Distinguishing normal brain aging from the development of alzheimer’s disease: inflammation, insulin signaling and cognition. J Neural regeneration research 13, 1719.

Dosenbach, N.U., Nardos, B., Cohen, A.L., Fair, D.A., Power, J.D., Church, J.A., Nelson, S.M., Wig, G.S., Vogel, A.C., Lessov-Schlaggar, C.N., 2010. Prediction of individual brain maturity using fmri. Science 329, 1358–1361.

Fjell, A.M., Westlye, L.T., Grydeland, H., Amlien, I., Espeseth, T., Reinvang, I., Raz, N., Holland, D., Dale, A.M., Walhovd, K.B., 2013. Critical ages in the life course of the adult brain: nonlinear subcortical aging. Neurobiol Aging 34, 2239–47. doi:10.1016/j.neurobiolaging.2013.04.006.

Franke, K., Dahnke, R., Clarke, G., Kuo, A., Li, C., Nathanielsz, P., Schwab, M., Gaser, C., 2016. Mri based biomarker for brain aging in rodents and non-human primates, in: 2016 international workshop on pattern recognition in neuroimaging (PRNI), IEEE. pp. 1–4.

Franke, K., Gaser, C., 2019. Ten years of brainage as a neuroimaging biomarker of brain aging: What insights have we gained? Frontiers in neurology 10, 789.

Franke, K., Gaser, C., Manor, B., Novak, V., 2013. Advanced brainage in older adults with type 2 diabetes mellitus. Frontiers in aging neuroscience 5, 90.

Franke, K., Ziegler, G., Klöppel, S., Gaser, C., Initiative, A.D.N., 2010. Estimating the age of healthy subjects from t1-weighted mri scans using kernel methods: exploring the influence of various parameters. Neuroimage 50, 883–892.

Gaser, C., Franke, K., Klöppel, S., Koutsouleris, N., Sauer, H., Initiative, A.D.N., 2013. Brainage in mild cognitive impaired patients: predicting the conversion to alzheimer’s disease. PloS one 8.

Good, C.D., Johnsrude, I.S., Ashburner, J., Henson, R.N.A., Friston, K.J., Frackowiak, R.S.J., 2001. A voxel-based morphometric study of ageing in 465 normal adult human brains. NeuroImage 14, 21–36. doi:https://doi.org/10.1006/nimg.2001.0786.

He, W., Goodkind, D., Kowal, P.R., 2016. An aging world: 2015.

Hötting, K., Röder, B., 2013. Beneficial effects of physical exercise on neuroplasticity and cognition. Neuroscience Biobehavioral Reviews 37, 2243–2257.

Johnson, T.E., 2006. Recent results: biomarkers of aging. Experimental gerontology 41, 1243–1246.

Jónsson, B.A., Bjornsdottir, G., Thorgeirsson, T., Ellingsen, L.M., Walters, G.B., Gudbjartsson, D., Stefansson, H., Stefansson, K., Ulfarsson, M., 2019. Brain age prediction using deep learning uncovers associated sequence variants. Nature communications 10, 1–10.

Lee, T., Sachdev, P., 2014. The contributions of twin studies to the understanding of brain ageing and neurocognitive disorders. Current opinion in psychiatry 27, 122–127.

Liang, H., Zhang, F., Niu, X., 2019. Investigating systematic bias in brain age estimation with application to post-traumatic stress disorders. Human Brain Mapping 40, 3143–3152. doi:https://doi.org/10.1002/hbm.24588.

Lu, T., Pan, Y., Kao, S.Y., Li, C., Kohane, I., Chan, J., Yankner, B.A., 2004. Gene regulation and dna damage in the ageing human brain. Nature 429, 883–891.

Löwe, L.C., Gaser, C., Franke, K., Initiative, A.D.N., 2016. The effect of the apoe genotype on individual brainage in normal aging, mild cognitive impairment, and alzheimer’s disease. PloS one 11.

MacNicol, E., Ciric, R., Kim, E., DiCenso, D., Cash, D., Poldrack, R., Esteban, O., 2020. Atlas-based brain extraction is robust across rat mri studies. [PrePrint] doi:10.31219/osf.io/sqxef.

MacNicol, E., Randall, K., Simmons, C., Kim, E., Turkheimer, F., Cash, D., 2019. Multimodal mr imaging of environmentally enriched and diet restricted rat model of healthy ageing, in: 2019 Neuroscience Meeting Planner. Chicago, IL: Society for Neuroscience.

MacNicol, E., Wright, P., Kim, E., Brusini, I., Esteban, O., Simmons, C., Turkheimer, F., Cash, D., 2021. Age-specific adult rat brain mri templates and tissue probability maps. [PrePrint] doi:10.31219/osf.io/htgqn.

Maioli, S., Puerta, E., Merino-Serrais, P., Fusari, L., Gil-Bea, F., Rimondini, R., Cedazo-Minguez, A., 2012. Combination of apolipoprotein e4 and high carbohydrate diet reduces hippocampal bdnf and arc levels and impairs memory in young mice. J Alzheimers Dis 32, 341–55. doi:10.3233/jad-2012-120697.

Martin, B., Mattson, M.P., Maudsley, S., 2006. Caloric restriction and intermittent fasting: two potential diets for successful brain aging. Ageing research reviews 5, 332–353.

Mattson, M.P., 2010. The impact of dietary energy intake on cognitive aging. Frontiers in aging neuroscience 2, 5.

Nenadić, I., Dietzek, M., Langbein, K., Sauer, H., Gaser, C., 2017. Brainage score indicates accelerated brain aging in schizophrenia, but not bipolar disorder. Psychiatry Research: Neuroimaging 266, 86–89.

Ngandu, T., Lehtisalo, J., Solomon, A., Levälahti, E., Ahtiluoto, S., Antikainen, R., Bäckman, L., Hänninen, T., Jula, A., Laatikainen, T., Lindström, J., Mangialasche, F., Paajanen, T., Pajala, S., Peltonen, M., Rauramaa, R., Stigsdotter-Neely, A., Strandberg, T., Tuomilehto, J., Soininen, H., Kivipelto, M., 2015. A 2 year multidomain intervention of diet, exercise, cognitive training, and vascular risk monitoring versus control to prevent cognitive decline in at-risk elderly people (finger): a randomised controlled trial. Lancet 385, 2255–63. doi:10.1016/s0140-6736(15)60461-5.

Pardoe, H.R., Cole, J.H., Blackmon, K., Thesen, T., Kuzniecky, R., Investigators, H.E.P., 2017. Structural brain changes in medically refractory focal epilepsy resemble premature brain aging. Epilepsy research 133, 28–32.

Pardoe, H.R., Kuzniecky, R., 2018. Napr: a cloud-based framework for neuroanatomical age prediction. Neuroinformatics 16, 43–49. doi:10.1007/s12021-017-9346-9.

Pedregosa, F., Varoquaux, G., Gramfort, A., Michel, V., Thirion, B., Grisel, O., Blondel, M., Prettenhofer, P., Weiss, R., Dubourg, V., 2011. Scikit-learn: Machine learning in python. the Journal of machine Learning research 12, 2825–2830.

Peters, R., 2006. Ageing and the brain. Postgraduate medical journal 82, 84–88.

Pfefferbaum, A., Rohlfing, T., Rosenbloom, M.J., Chu, W., Colrain, I.M., Sullivan, E.V., 2013. Variation in longitudinal trajectories of regional brain volumes of healthy men and women (ages 10 to 85 years) measured with atlas-based parcellation of mri. Neuroimage 65, 176–93. doi:10.1016/j.neuroimage.2012.10.008.

Quinn, R., 2005. Comparing rat’s to human’s age: how old is my rat in people years? Nutrition 21, 775.

Rando, T.A., Chang, H.Y., 2012. Aging, rejuvenation, and epigenetic reprogramming: resetting the aging clock. Cell 148, 46–57.

Raz, N., Lindenberger, U., Rodrigue, K.M., Kennedy, K.M., Head, D., Williamson, A., Dahle, C., Gerstorf, D., Acker, J.D., 2005. Regional brain changes in aging healthy adults: general trends, individual differences and modifiers. J Cerebral cortex 15, 1676–1689.

Robinson, S.D., Dymerska, B., Bogner, W., Barth, M., Zaric, O., Goluch, S., Grabner, G., Deligianni, X., Bieri, O., Trattnig, S., 2017. Combining phase images from array coils using a short echo time reference scan (composer). Magnetic resonance in medicine 77, 318–327.

Ronan, L., Alexander-Bloch, A.F., Wagstyl, K., Farooqi, S., Brayne, C., Tyler, L.K., Fletcher, P.C., 2016. Obesity associated with increased brain age from midlife. Neurobiology of aging 47, 63–70.

Soininen, H., Solomon, A., Visser, P.J., Hendrix, S.B., Blennow, K., Kivipelto, M., Hartmann, T., group, t.L.c.s., 2021. 36-month lipididiet multinutrient clinical trial in prodromal alzheimer’s disease. Alzheimer’s Dementia 17, 29–40. doi:https://doi.org/10.1002/alz.12172.

Speisman, R.B., Kumar, A., Rani, A., Pastoriza, J.M., Severance, J.E., Foster, T.C., Ormerod, B.K., 2013. Environmental enrichment restores neurogenesis and rapid acquisition in aged rats. Neurobiology of aging 34, 263–274.

Steffener, J., Habeck, C., O’Shea, D., Razlighi, Q., Bherer, L., Stern, Y., 2016. Differences between chronological and brain age are related to education and self-reported physical activity. Neurobiology of aging 40, 138–144.

Steiger, J.H., 1980. Tests for comparing elements of a correlation matrix. Psychological Bulletin 87, 245–251. doi:10.1037/0033-2909.87.2.245.

Teter, B., Finch, C.E., 2004. Caliban’s heritance and the genetics of neuronal aging. Trends in neurosciences 27, 627–632.

Turkheimer, F.E., Rosas, F.E., Dipasquale, O., Martins, D., Fagerholm, E.D., Expert, P., Váša, F., Lord, L.D., Leech, R., 2021. A complex systems perspective on neuroimaging studies of behavior and its disorders. Neuroscientist, 1073858421994784doi:10.1177/1073858421994784.

Vinke, E.J., de Groot, M., Venkatraghavan, V., Klein, S., Niessen, W.J., Ikram, M.A., Vernooij, M.W., 2018. Trajectories of imaging markers in brain aging: the rotterdam study. Neurobiology of Aging 71, 32–40. doi:https://doi.org/10.1016/j.neurobiolaging.2018.07.001.

Walhovd, K.B., Fjell, A.M., Reinvang, I., Lundervold, A., Dale, A.M., Eilertsen, D.E., Quinn, B.T., Salat, D., Makris, N., Fischl, B., 2005. Effects of age on volumes of cortex, white matter and subcortical structures. Neurobiology of Aging 26, 1261–1270. doi:https://doi.org/10.1016/j.neurobiolaging.2005.05.020.

Wood, T.C., Simmons, C., Hurley, S.A., Vernon, A.C., Torres, J., Dell’Acqua, F., Williams, S.C., Cash, D., 2016. Whole-brain ex-vivo quantitative mri of the cuprizone mouse model. PeerJ 4, e2632.

Yorke, A., Kane, A.E., Hancock Friesen, C.L.,Howlett, S.E., O’Blenes, S., 2017. Development of a rat clinical frailty index. The Journals of Gerontology: Series A 72, 897–903.

