## Supplementary Material for "MRI-derived brain age as a biomarker of ageing in rats: validation using a healthy lifestyle intervention"

Table A1: Comparison of the mean absolute error (MAE) obtained by changing the tissue probability maps (TPMs) used as input to the GPR model. These results are obtained using leave-one-out cross-validation on the training cohort. The best performing model, which was then employed for the present study, is indicated in bold.

| **Input type** | | |
|---|---|---|
| **TPMs** | **Templates** | **MAE (weeks)** |
| GM | 11 months | 5.84 |
| WM | 11 months | 6.53 |
| GM + WM | 11 months | 5.76 |
| **GM + WM + CSF** | **11 months** | **5.69** |
| GM | All | 5.80 |
| WM | All | 6.53 |
| GM + WM | All | 5.79 |
| GM + WM + CSF | All | 5.72 |
